# Lack of dcf1 leads to neuronal migration delay, axonal swollen and autism-related deficits

**DOI:** 10.1101/2020.02.20.958934

**Authors:** Ruili Feng, Yanlu Chen, Yangyang Sun, Guanghong Luo, Jianjian Guo, Qiang Liu, Jie Wu, Xiangchun Ju, Tieqiao Wen

## Abstract

Perturbed neuronal migration and abnormal axonogenesis have been shown to be implicated in the pathogenesis of autism spectrum disorder (ASD). However, the molecular mechanism remains unknown. Here we demonstrate that dendritic cell factor 1(DCF1) is involved in neuronal migration and axonogenesis. The deletion of *dcf1* in mice delays the localization of callosal projection neurons, while dcf1 overexpression restores normal migration. Delayed neurons appear as axon swelling and axonal boutons loss, resulting in a permanent deficit in the callosal projections. Western blot analysis indicates that absence of dcf1 leads to the abnormal activation of ERK signal. Differential protein expression assay shows that PEBP1, a negative regulator of the ERK signal, is significant downregulation in *dcf1* KO mice. Direct interaction between DCF1 and PEBP1 is confirmed by Co-immunoprecipitation test, thus indicating that DCF1 regulates the ERK signal in a PEBP1-dependent pattern. As a result of the neurodevelopmental migration disorder, *dcf1* deletion results in ASD-like behaviors in mice. This finding identifies a link between abnormal activated ERK signaling, delayed neuronal migration and autistic-like behaviors in humans.

## Introduction

The development of the cerebral cortex is a complicated process involving cell differentiation, migration, axonogenesis and formation of neural circuits (Borrell & Reillo, 2012). Studies have shown that cortical projection neurons in the developing brain are generated within the ventricular (VZ) and subventricular (SVZ) zones in the dorsal telencephalon(Franco & Muller, 2013; LaMonica et al, 2012) and migrate in an inside-out fashion along radial trajectories to reach their destiny, where they eventually form neuronal circuits(Sur & Leamey, 2001). Timing of migration and localization is crucial for the proper integration of neurons into cortical circuits (Bocchi et al, 2017; He et al, 2015; Sawada et al, 2018). Callosal projection neurons (CPNs) mainly produced in E15 with medium pyramidal size that are primarily located in layers II/V, and extend an axon across the corpus callosum (CC) to distant intracortical, subcortical and subcerebral targets (Molyneaux et al, 2007). Delayed migration of callosal projection neurons without apparent structural defects may impact on the subsequent development of cortical circuits and cause altered interhemispheric connections and impaired social behavior(Bocchi et al, 2017; Mata et al, 2009; Yeung et al, 1999). Despite some progress in this respect, there is always a lack of understanding the link between transient migration delays and abnormal social behaviors.

ERK is the most downstream protein kinase of the canonical Ras-Raf-MEK-ERK cascade that can be suppressed by Raf kinase inhibitory protein, PEBP1(Pyo et al, 2018). ERK signaling is critical in early neurogenesis (Ye et al, 2016); it directs self-renewal, division and differentiation of neural progenitor cells(Dee et al, 2016; Pillat et al, 2016). ERK signaling is dispensable for morphological and survival of large Ctip2^+^ neurons in layer 5(Xing et al, 2016). RASopathy patients with neurodevelopmental delay, cognitive impairment, and epilepsy are most often associated with hyper-active ERK/MAPK signaling (Nowaczyk et al, 2014). Little is known about how ERK/MAPK signaling might relate to the migration of projection neurons during early neural development and pathogenesis of autism.

DCF1, also referred to as TMEM 59, a membrane-bound protein, is involved in neural stem cells (NSCs) in differentiation and morphogenesis(Wang et al, 2008; Wen et al, 2002). Silencing dcf1 inhibited the proliferation of NSCs in vitro. Knockout of dcf1 leads to dendritic spine dysplasias, which in turn causes of the damage to learning and memory (Liu et al, 2018). DCF1 promotes the formation of multi-component Wnt-FZD assembly through intra-membrane interactions. These Wnt-fzd-DCF1 clusters then merged with LRP6 to form mature Wnt signalosomes (Gerlach et al, 2018). Here we asked how loss of dcf1 hinders neuronal migration and identified new dcf1-mediated signal. We discovered an extraordinary role of dcf1 in regulating the radial migration and axonal morphogenesis of projection neurons via ERK signal. Excessive activation of ERK signals caused by DCF1 knockout induced a short delay in the radial migration of CPNs in the somatosensory cortex of mice by reducing the speed of neuronal movement. Overexpression of DCF1 / PEBP1 or administration of ERK inhibitor restored normal migration. This supports an unexpected link of canonical ERK signaling, neuronal migration, swollen axons, and ASD-like behaviors.

## Results

### DCF1 is essential for proper lamination of the cerebral cortex

As previously mentioned (Liu et al, 2018), *dcf1-/-* mice were hybridized with Thy1-GFP line M mice. Thy1-positive neurons labeled with GFP should be located in layer 5 in primary somatosensory area in Thy1-GFP; *dcf1+*/+ mice, but they were anomalously present at layer 2 to 3 in Thy1-GFP; *dcf1*-/- mice (Fig. 1A. indicated by red arrow). To confirm the abnormal lamination of primary somatosensory cortex due to dcf1 knockout, we immunostained coronal sections of the adult brain using antibodies to the neuronal marker NEUN (Fig. 1B). Further analysis revealed that dcf1 KO mice had thinner cortices and significantly fewer neurons than wild-type mice (counting NeuN^+^ cells within 200μm in the barrel cortex). In DCF1 KO mice, the cortex was divided into 10 equal parts, and the number of neurons was significantly reduced in the superficial cortex and increased significantly in the deep cortex (Fig. 1C). Similarly, western blot results indicated significant reduction in neuron-specific marker, as well as in astrocyte and oligodendrocyte markers (Fig. 1 D and E).

**Fig 1.**
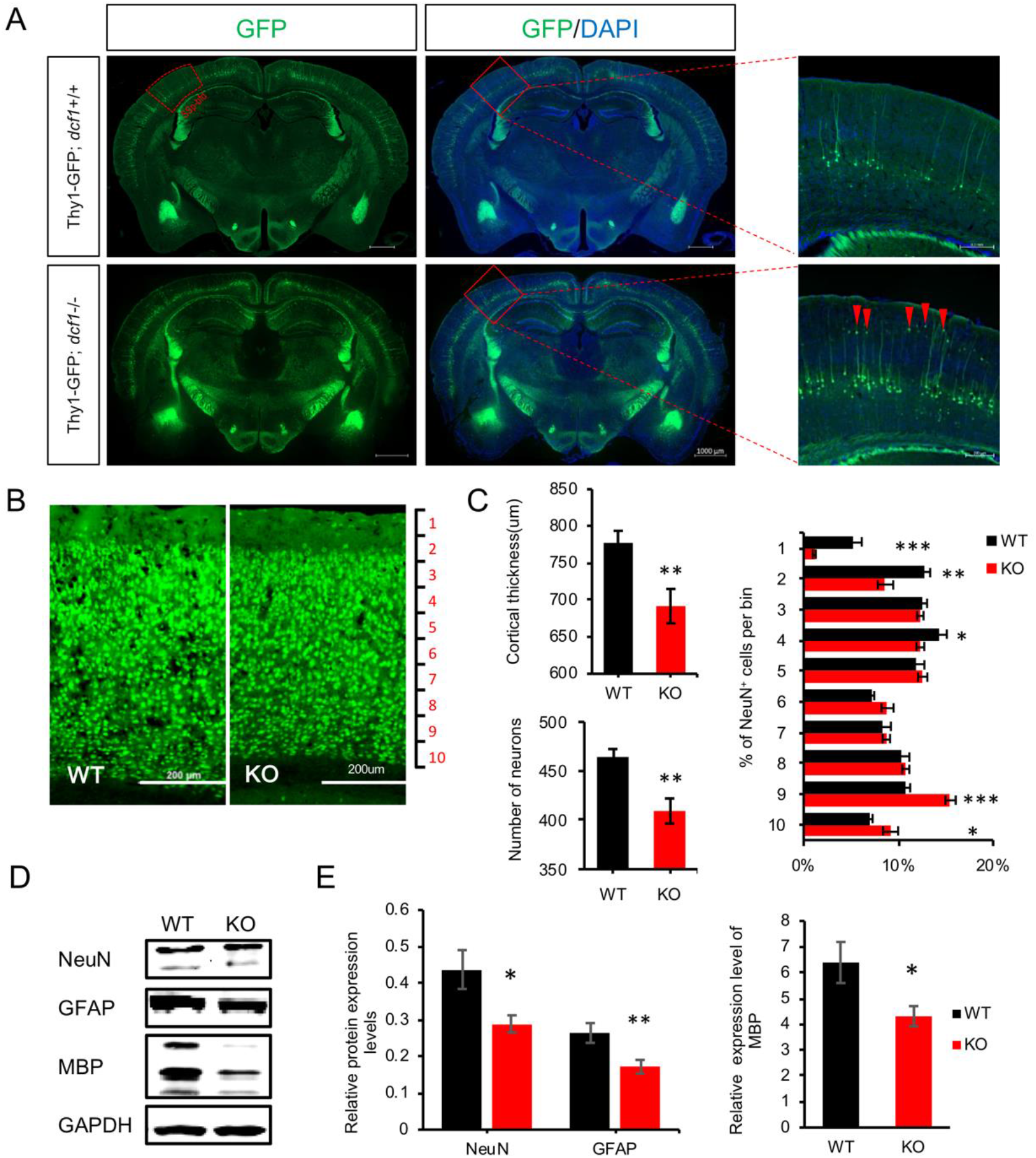
Abnormal location of cortical cells in *dcf1*-null mice. (A) *dcf1-/-* mice were crossed with Thy1-GFP line M mice. Thy1-positive neurons were labeled with GFP. GFP-positive neurons were located in layer 5 of the neocortex, whereas the deletion of *dcf1* cause some neurons mislocate at the 2-3 layers in the barrel cortex of Thy1-GFP; *dcf1*-/- mice (indicated by red arrow). (B) Coronal sections of the cerebral cortex were immunostained with anti-NEUN. (C) Thinner cortical thickness and fewer neurons were identified in the neocortex of *dcf1* KO mice. Further, percentage of NeuN labeled cells across 10 equal bins from pia to white matter were calculated at the width of 200 µm in *dcf1* KO mice. Neurons located at superficial layers were significantly decreased, whereas neurons at deeper layers were substantially increased. (4 mice in each group, 3 consecutive sections per mouse, Student’s t-test, *p < 0.05, **p < 0.01). (D, E) Western blotting was also indicated a reduction of NEUN, GFAP and MBP expressions in *dcf1* KO mice cerebral cortex. (Student’s t-test, n≥4, *p < 0.05, **p < 0.01, ***p < 0.001). (SSp-bfd, Primary somatosensory area, barrel field).

### Dcf1 ablation impairs neuronal migration and axonal morphogenesis

Given the abnormal lamination of the cortex in the dcf1 KO mice, we thus pursued an investigation into the migration of cortical neurons. At E17.5 the cortical neurons in the control group were tightly clustered in a radial pattern, but were disorganized in the dcf1-null mice (Fig S1A indicated by arrows). Subsequently, in utero electroporation was performed to investigate neural migration. The experimental pattern and schedule are shown in figure 2A and B. An EGFP-encoding plasmid was electrotransfered into cortical progenitor cells in the ventricular zone of the embryonic brain at E14.5. The callosal projection neurons (CPNs), which were mainly produced in E15, were EGFP-positive and had a medium pyramidal size, destined for layer IV. When sliced at E17.5, neuronal migration pattern of EGFP-marked cells could be observed in the neocortex in the dorsal lateral area. In dcf1 KO mice, we observed a decrease in the distribution of EGFP^+^ neurons in the surface layer of cortex (up cortex plate) and an increase distribution of the middle and low cortex plate (Fig. 2 C and D), indicating a significant difference in the migration of CPNs between the WT and dcf1-/- mice. Similar results were visualized in newborn neurons via immunohistochemical analysis following bromodeoxyuridine (BrdU) administration (Fig. S2). At P21, we observed that EGFP^+^ neurons were mainly destined in Layer4 in wild-type mouse, which is completely different from dcf1 KO mice mainly distributed in shallower Layer2/3 (Fig.2E). Immunofluorescence staining of cux1, a marker of neurons in layers 2-4, further demonstrated the delay of neuronal localization caused by DCF1 knockout (Fig. 2F). The deletion of dcf1 not only affects the migration of neurons, but also affects the morphology of neuronal axon. Loss of boutons and swollen of axons were observed in prenatal and postnatal *dcf1-/-* mice (Fig. 2G and H). Frozen section at E17.5 revealed that the intermediate zone (IZ) neurons in wild type mice had finer axons and rich boutons. By contrast, the neuronal axon in dcf1-null mice was swollen, and boutons didn’t appear (Fig. 2G). At P21, axons could be observed undisturbed in the corpus callosum, by which the CPNs extend axons to distant targets. The axons in the corpus callosum in DCF1 KO mice remained swollen, although the button and its size have been significantly reduced (Fig. 2H). In general, dcf1 knockout affects nerve development, characterized by delayed nerve migration, axonal swelling, and abnormal boutons.

**Fig 2.**
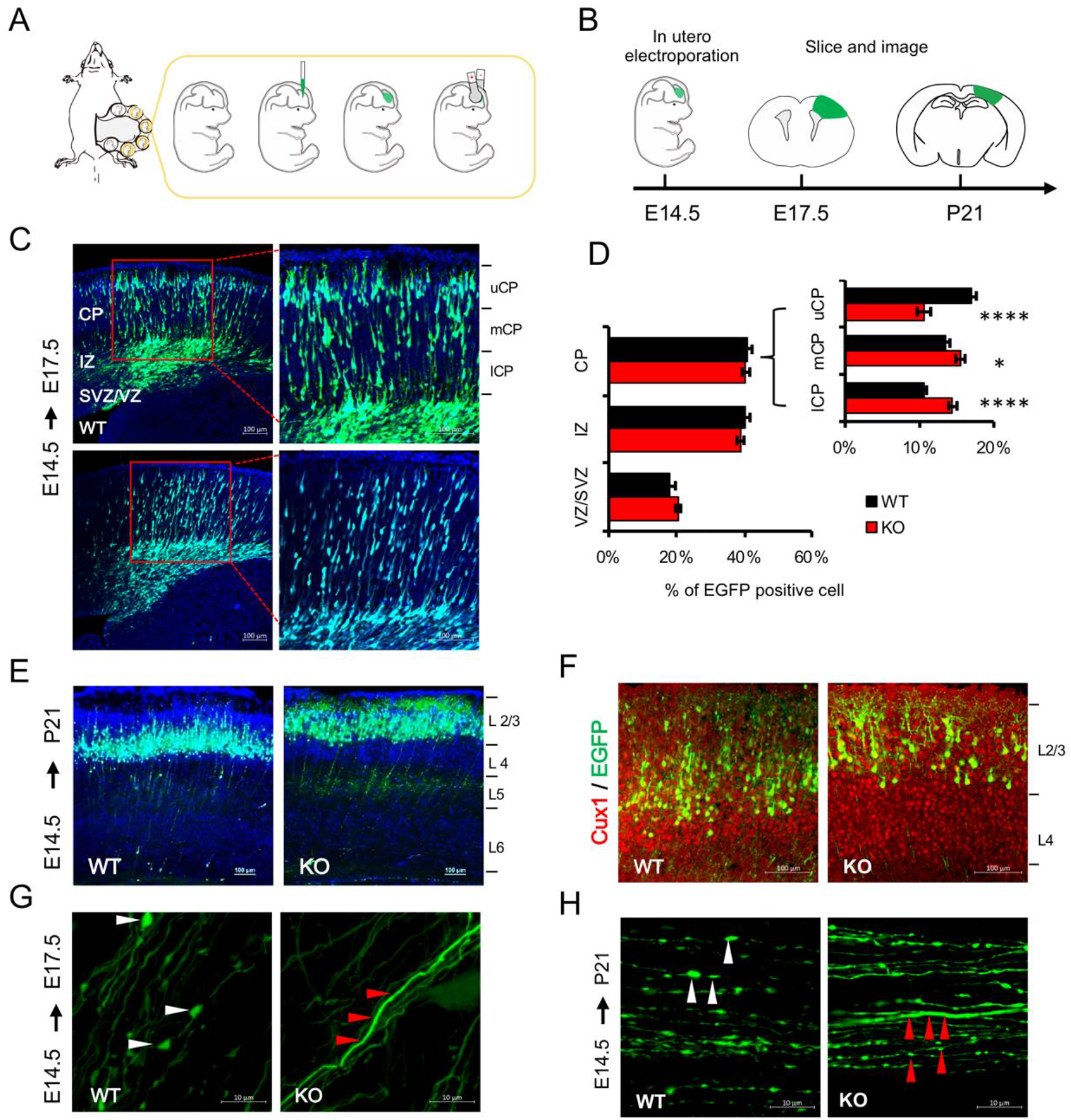
Dysplasia of cortical neurons in the absence of dcf1. (A). Experimental pattern diagram of In utero electroporation. (B) Experimental schedule. At E14.5, cortical progenitor cells in the ventricular zone of the mouse brain were transfected with pCAGGS-EGFP via in utero electroporation and were sectioned at E17.5 or P21 to observe neural migration in neocortex. (C) By DAPI marker, the cortical wall could be divided into the cortical plate (CP), intermediate zone (IZ) and subventricular zone/ventricular zone (SVZ/VZ). CP was further subdivided into upper CP (uCP), median CP (mCP) and lower CP (lCP). (D) The graph presented the quantification of neuronal migration in *dcf1* KO mice and control. The fraction of EGFP^+^ cells reached different zones of the cortex were measured at 3 days after electroporation. Data were presented as the mean ± sem from 12 sections prepared from three or four embryos obtained from two or three littermates. (Student’s t test; *p < 0.05, ****p<0.0001.) (E) When sectioned at P21, cortical neurons were mislocated at layer II-III in KO mice, however, wild-type neurons were destined for layer II-IV. (F) Immunofluorescence staining of Cux1, a marker of neurons in layer II-IV. In the control mice, EGFP^+^ neurons had arrived at layer II-IV, whereas in the *dcf1-/-* mouse, most EGFP-expressing cells were dislocated at the top layer II-III. (G) EGFP^+^ neurons in WT cortex possessed many axonal boutons (white arrows) in the intermediate zone (IZ) at E17.5, whereas loss of boutons and swollen of axons were observed in *dcf1-/-* mice (red arrows). (H) when observed at P21, these EGFP^+^ cells extended long-distance axons through the corpus callosum and projected to the contralateral hemisphere, axons of wild-type mice possessed many larger boutons in corpus callosum. However, boutons of *dcf1-/-* mice were smaller and axons were swollen (red arrows).

### Neurons with delayed migration exhibit slower velocity in dcf1-null mice

To find out the causes of the abnormal localization of neurons in dcf1 KO mice, the electroporated cortex (E17.5) was subjected to vibrational sections and continuously cultured for 4 hours on the Live Cell Imaging System, and the process of neuronal migration was recorded by time-lapse imaging (Fig. 3 A-a, b). After tracing the living cells with Imaris, it was found that neurons migrated significantly slower in KO mice than in WT mice (P< 0.01) (Fig. 3 A-a, b and d, Supplementary Material: Videos S1 and S2). DCF1 re-expression could accelerate neuronal migration (Fig. 3 A-c and d, Supplementary Material: Videos S2 and S3), thereby rescuing for the migration and axons defects caused by dcf1 deletion (Fig. 3B, C).

**Fig 3.**
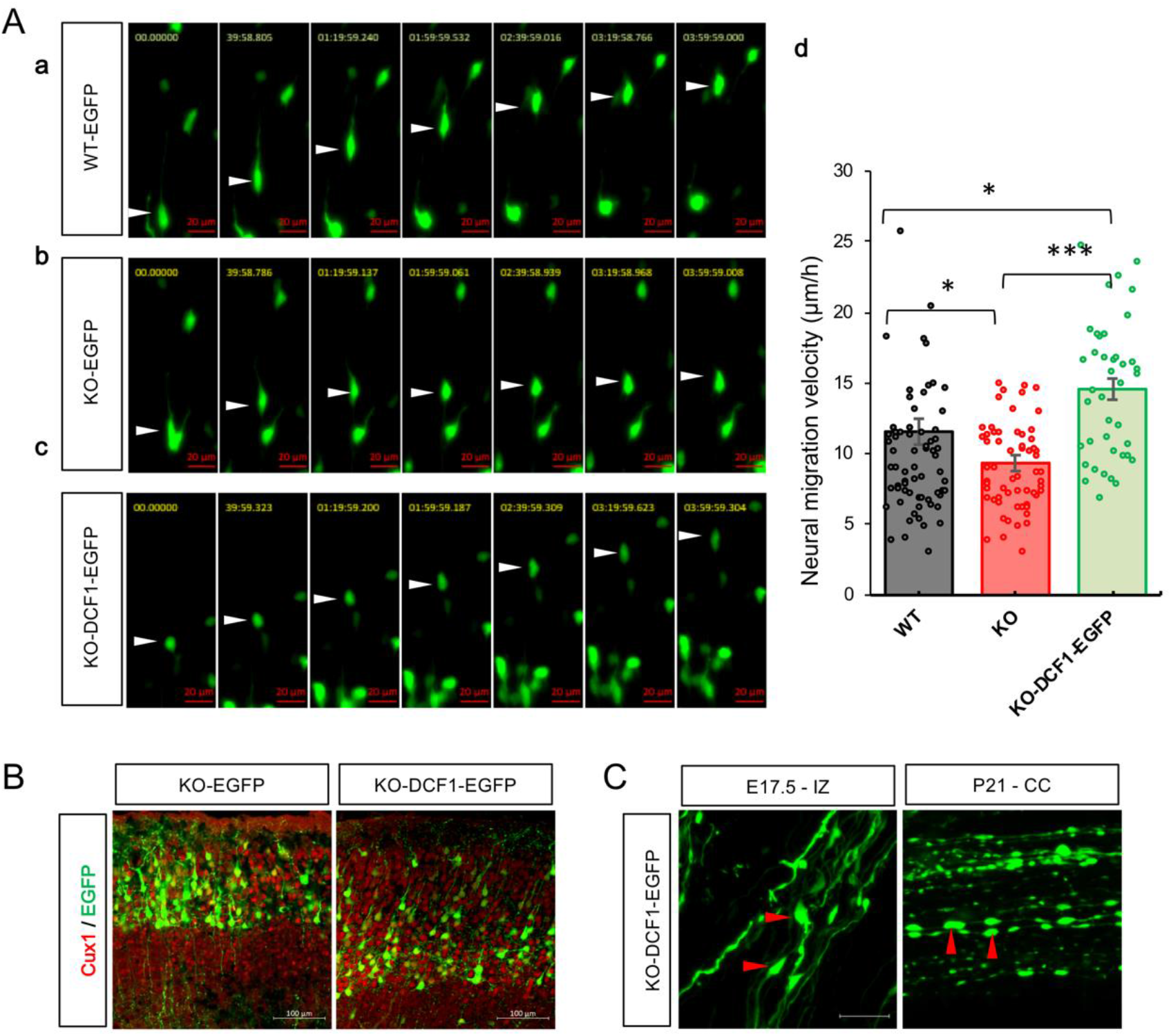
Deficits of neuronal migration and morphology in dcf1-/- mice can be rescued by re-expression of d*cf1*. (A) Embryonic electroporation was performed on pregnant mice at E14.5 days, and vibration slice of fetal brain was taken and cultured till E17.5. Continuous observation was performed on the living cell imaging station by time-lapse technique for 4 hours (take a photo every 5 minutes). (a) Embryonic brains in the wild type mice were electroporated with pCAGGS-EGFP; (b) Embryonic brains in *dcf1* KO mice were electroporated with pCAGGS-EGFP; (c) Embryonic brains in *dcf1* KO mice were rescued by transfering pCAGGS-DCF1-EGFP. From the comparison of their migration rates, the loss *dcf1* leads to the slowest migration. (d) Statistical analysis of migration velocity. Neuronal migration velocity (μm/h) in the *dcf1-/-* mice were significantly decreased compared with that of the controls and could be rescued by pCAGGS-DCF1-EGFP. (*p < 0.05, **p < 0.01, ***p < 0.001, n_WT_=70, n_KO_=69, n_KO-DCF1_=40). (B) At P21, anti-cux1 immunofluorescence assay showed that re-expression of DCF1 could help neurons locate in the cortex correctly, showing that EGFP positive neurons in the KO mice brains had distribute over layer II-IV.(C) Swollen of axons and loss of boutons were repaired by re-expression of dcf1 in KO mice both at E17.5 and P7 (red arrows).

### Dcf1 deficiency stunts neuronal migration by ERK signaling pathway

It is generally believed that JNK, one of the MAPK signaling pathway, contributes to neuronal migration(Huang et al, 2004; Tarcic & Yarden, 2010; Yamasaki et al, 2012), so the phosphorylation level of JNK was detected by western blot to evaluate its activity. After standardization to GAPDH, the mean level of JNK and phosphorylated JNK in the cortex of dcf1 KO mice at E14.5, E17.5, P1 and P7 did not change compared to their WT littermates (Fig. 4 A, B). Surprisingly, the ERK1/2 signaling in dcf1 KO mice was significantly activated at different developmental periods (Fig. 4 A, B). Further, the functional complementation analysis indicated that the overexpression of DCF1 in HEK 293T cells significantly decreased the p-ERK1/2 levels (P< 0.01) (Fig. 4 C), even in the presence of PMA, an activator of ERK signals (P< 0.01) (Fig. 4 D). Strikingly, the activation of ERK signaling was rescued by treatment of a MEK/ERK inhibitor (PD98059) during embryonic development stages (Fig. 4 E). Therefore, DCF1 acts on the ERK signaling pathway by affecting the proteins located in the upstream of MEK and downstream of PKC. We argued whether reducing p-ERK level in dcf1-/- neurons could correct their migration defects. It was confirmed by intraperitoneal injection of PD98059 in dcf1 KO mice. Results indicated that inhibition of ERK signal could repair the neuronal migration anomaly from VZ/SVZ to IZ caused by dcf1 deletion (Fig. 4 F and G). Taken together, these results support the idea that the absence of dcf1 leads to the abnormal activation of ERK signal, resulting in delayed neuronal migration.

**Fig 4.**
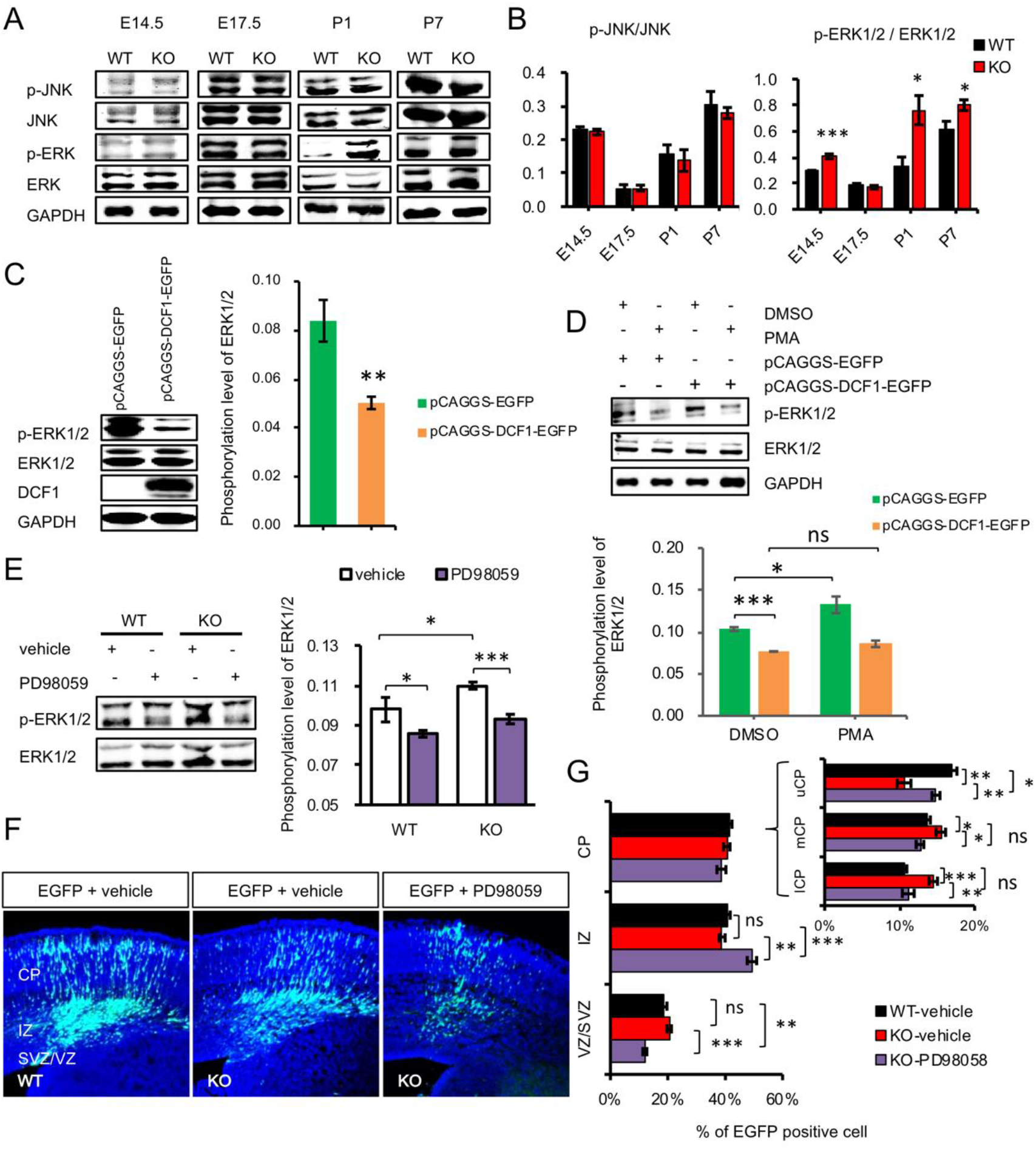
*Dcf1* affects cortical neural development through ERK signaling pathway. (A) Western blot was used to analyze activity of the JNK and ERK signaling pathway at E14.5, E17.5, P1 and P7 in WT and *dcf1*-null mice. (B) Graph representing the quantitative analysis of phosphorylated JNK and ERK. (C) HEK293T cells were transfected with pCAGGS-EGFP and pCAGGS-DCF1-EGFP, and the effect of overexpression of DCF1 on ERK phosphorylation was analyzed (D) 24 h after transfection, PMA (activator of ERK pathway) was added to the culture medium and the change of phosphorylated ERK was analyzed by electrophoresis. (E) Heterozygous self-mating pregnant female mice were administered PD98059 (10mg/kg, ip qd), an inhibitor of MEK and ERK, from E14.5 to E17.5. At E17.5, cortical proteins were extracted from fetal mice and the changes of activated ERK were detected in WT and *dcf1* KO mice. (F, G) Pregnant mice electroporated with plasmid EGFP at E14.5 were administered PD98059 from E14.5 to E17.5. Representative images of EGFP^+^ neurons (F) and statistical analysis of cortical migration (G). Data were shown as Mean ± SEM; n>4, Student’s t-test; *p < 0.05, **p < 0.01, ***p < 0.001.

### DCF1 may act on the ERK pathway through PEBP1

To find possible regulatory factors for ERK signaling in the absence of dcf1, we performed a differential protein expression assay and found PEBP1, a negative regulator of the ERK signal, was significant downregulation in adult dcf1 KO mice (FIG S3). To study the expression pattern of PEBP1 in neurodevelopmental stage, we used western blot to detect the relative expression level of PEBP1 at multiple time points. Results showed that PEBP1 expression in dcf1-null mice was reduced at E14.5, E17.5 and P1 (Fig. 5 A). Since PEBP1 is an inhibitor of the ERK signal, this phenomenon was matched with the hyperactivation of ERK, although the expression of PEBP1 at P7 was increased due to feedback regulation (Fig. 5 A). Functional complementation analysis indicated that overexpression of DCF1 in HEK293 cells could significantly upregulate the level of PEBP1 (Fig. 5 B). The combination of DCF1 and PEBP1 was investigated by isothermal titration calorimetry (ITC) with a Kd of 6.44E5 ± 1.14E5 M^-1^, indicating a high binding affinity of DCF1 to PEBP1 (Fig. 5 C). The direct interaction between DCF1 and PEBP1 was also confirmed by co-immunoprecipitation experiment (Fig. 5 D).

**Fig 5.**
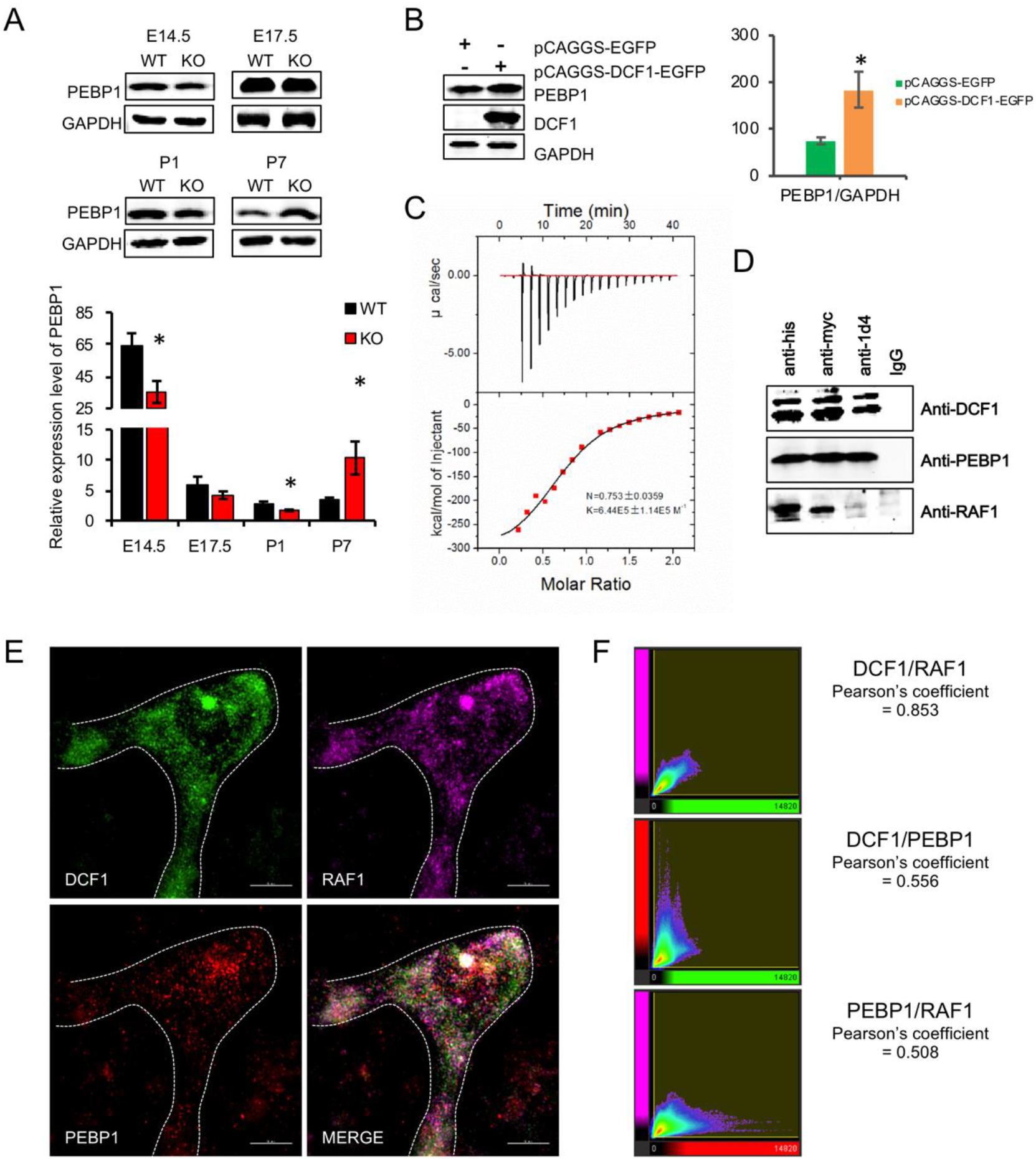
PEBP1 and RAF1 may be involved in the functioning of DCF1. (A) Representative images of PEBP1 expression at E14.5, E17.5, P1 and P7 in *dcf1* KO mice and WT controls (Data was shown as Mean ± SEM; n=4, Student’s t-test; *p < 0.05, **p < 0.01, ***p < 0.001.). (B-D) Representative images of the interaction between DCF1 and PEBP1. (B) Western blot and statistical analysis of PEBP1 expression in HEK293T cells transfected with pCAGGS-EGFP and pCAGGS-DCF1-EGFP. (C) ITC titration analysis of DCF1 and PEBP1 interaction. Parameters obtained from this experiment are: N = 0.753 ± 0.0359; Kd = 6.44E5 ± 1.14E5 M^-1^; Δ*H* = −3.291E5 ± 2.247E5 cal/mol; ΔS = −1.08E3 cal/mol/deg. (D) HEK293T cells were transfected with pcDNA3.1-DCF1-his-myc or pcDNA3.1-PEBP1-1d4 respectively, lysed and subjected to co-immunoprecipitation with agarose beads coupled to anti-his, anti-myc and anti-1d4 antibodies. Immunoprecipitation products were detected using Western blotting with the indicated antibody. (E) After immunostaining of primary cultured cortical neurons for DCF1, PEBP1 and RAF1, SIM super-resolution imaging was performed and the Pearson correlation coefficient (PCC) analysis demonstrated that these proteins relative cellular co-localized with each other (PCC>0.5).

Given that PEBP1 inhibits the ERK signaling by binding RAF1 (Yeung et al, 1999), we used anti-RAF1 to detect the NC membrane of the above co-immunoprecipitation, confirming the interact between RAF1, DCF1 and PEBP1 (Fig. 5 D). To investigate whether these proteins interact with each other in neurons, primary cultured cortical neurons were imaged by SIM super-resolution microscopy with immunostaining (Fig. 5 E). Pearson correlation coefficient analysis showed there was a colocation between DCF1, PEBP1 and RAF1 (Fig. 5 F). Thus, the molecular mechanism of dcf1 regulating ERK signal was confirmed. Subsequently, we wondered whether over expression of PEBP1 could repair the damage caused by dcf1 knockout. Cortical progenitor cells were transfected with pCAGGS-PEBP1-EGFP at E14.5 and sliced at E17.5 (Fig. 6A), quantification of migration neurons shows that PEBP1 could reverse the effect of dcf1 knockout on neuronal migration (P< 0.01) (Fig.6B). At P21, with the expression of PEBP1, the repair of neuronal migration was clearly visible (Fig. 6 C). These results support our hypothesis that DCF1 regulates the ERK signal in a PEBP1-dependent pattern.

**Fig. 6.**
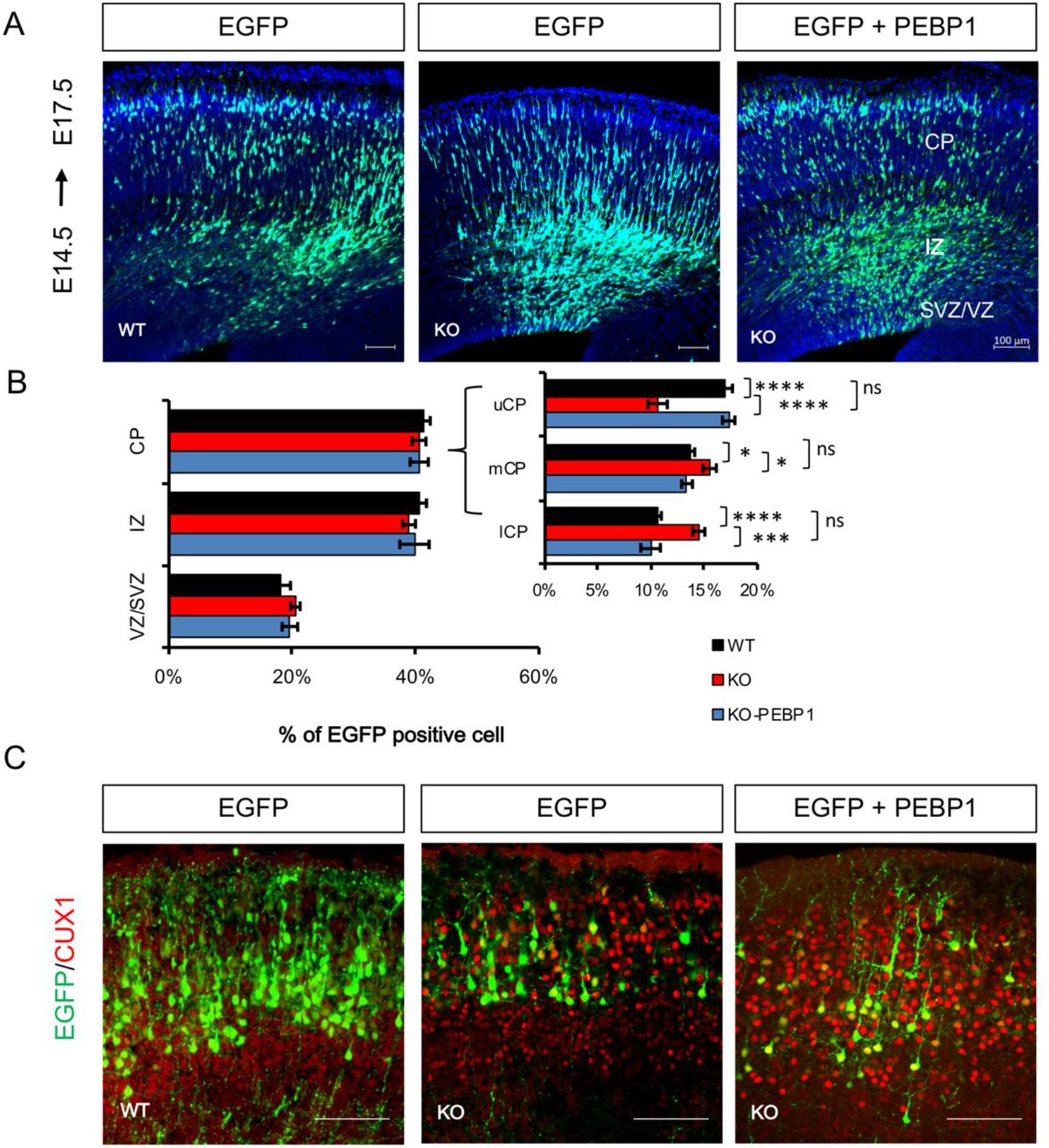
PEBP1 promotes radial migration in *dcf1-/-* mouse. (A) Cortical progenitor cells were transfected with pCAGGS-PEBP1-EGFP via in utero electroporation at E14.5 and sliced at E17.5. (B) Quantification of the migration neurons showed that PEBP1 could reverse the effects of knockdown of *dcf1* on neuronal migration. (C) At P21, the EGFP^+^ neurons rescued by PEBP1 overexpression in the KO mice mainly dispersed from layer II to IV.

### *Dcf1* deletion results in ASD-like behavior in mice

Recent studies have shown that transient migration delays without apparent structural defects may affect subsequent development of cortical circuits and produce autistic behaviors (Bocchi et al, 2017). Hyperactivation of ERK signal is also associated with autism (Courchesne et al, 2019; Faridar et al, 2014; Gazestani et al, 2019; Pinto et al, 2010). ASD is characterized by persistent difficulties in social communication and interaction, restricted interests and repetitive behaviors, as well as learning and memory deficits (Pasciuto et al, 2015). We employed a three-chamber assay to probe voluntary social interaction (Kaidanovich-Beilin et al, 2011). *Dcf1* KO mice and WT mice had no preference for the empty cage, but both *dcf1* KO mice and WT mice were more interested in a novel mouse than the inanimate empty cages (Fig. 7 A). And then we did the social novelty test by distinguishing between a familiar mouse and a new unfamiliar one. WT mice showed a strong preference for the compartment containing unfamiliar mouse (stranger 2), while *dcf1* KO mice failed to distinguish between the familiar (stranger 1) and unfamiliar mouse (Fig. 7 B). Furthermore, we performed a reciprocal interaction test. In different combinations of mice (WT-WT, WT-KO, KO-KO) and different environments (Home cage or New cage), the number of contacts and time between mice were counted. It was found that the close contact time for KO-KO group was significantly lower than that of WT-WT and WT-KO in familiar environment, and the contact times and time of KO-KO and WT-KO were significantly lower than that of WT-WT in unfamiliar environment, indicating that *dcf1* KO mice had decreased social interaction compared with the WT controls (Fig. 7 C). Furthermore, in the social dominant tube test, KO mice exhibited more dominant behavior than the controls (Fig. 7 D). We then examined whether KO mice exhibited repetitive and perseveration behaviors, which are behavioral basis for ASD diagnosis (Rapin & Tuchman, 2008). In the marble burying task, the number of buried marbles in *dcf1* KO mice was significantly more than that in WT mice, which indicated a more severe obsessive-like symptom in *dcf1* KO mice (Fig. 7 E). We also examined nest building behavior. In KO mice, the cotton pads in the cage were severely torn. The quality of the nest was poor, and more than 90% of mice had failed to build their nests, indicating that the *dcf1* KO mice were also significantly impaired during the task (Fig. 7 F). In the Y maze test, the cognitive plasticity in *dcf1* KO mice was defective (Fig. 7 G). We then performed open field test, which is thought to be related to ASD (Guo et al, 2018). *Dcf1* KO mice exhibited poor locomotion activity and increased self-grooming behavior (Fig. 7H). Taken together, these results demonstrate that *dcf1* deletion could cause ASD-like behavior in mice.

**Fig. 7.**
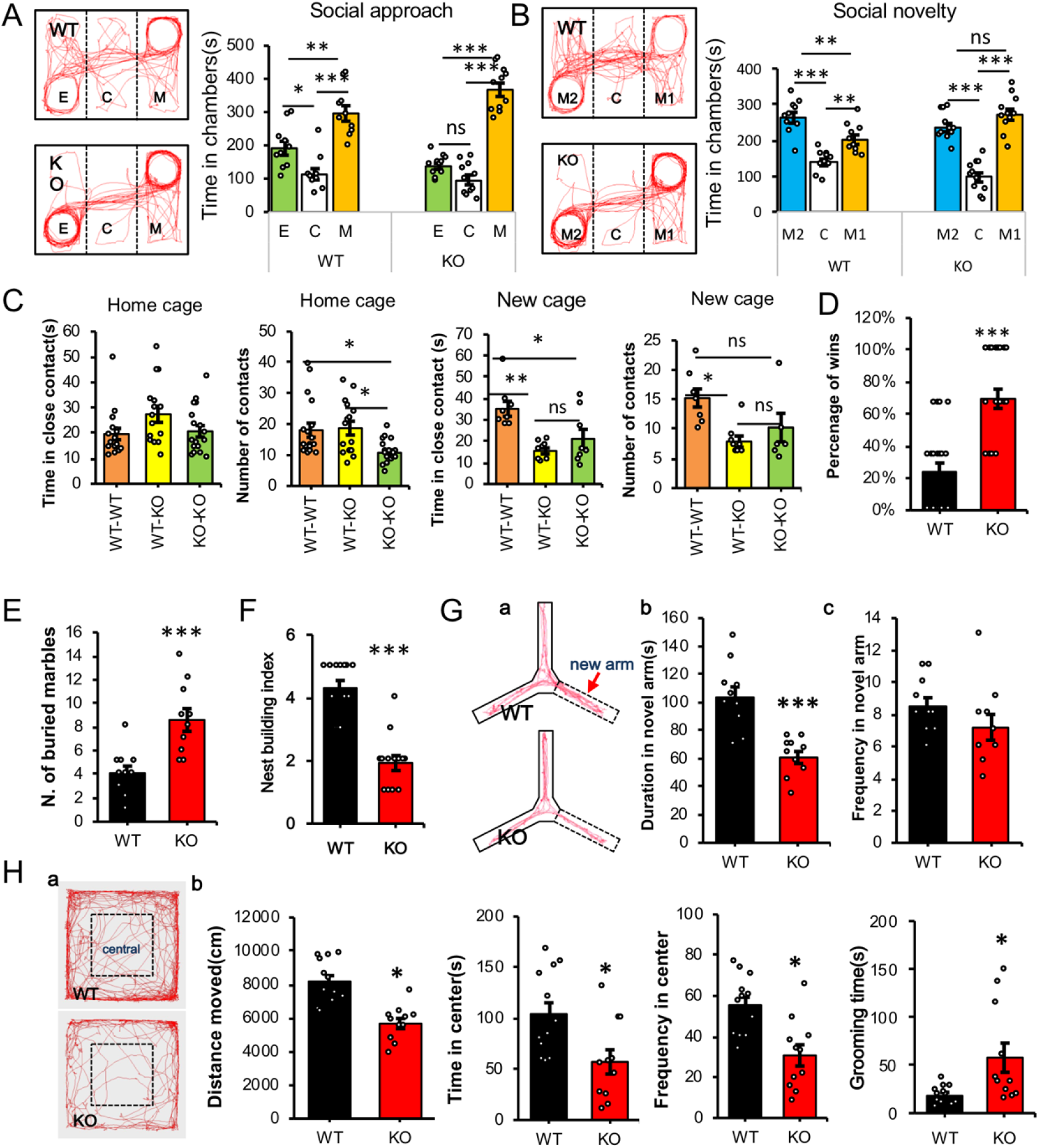
Loss of *dcf1* leads to ASD-like behaviors. (A, B) Three-chamber experiment was used to test social preference and social novelty. Both WT and KO mice have no preference for the left and right sides of a three-chamber unit during the habituation period. (A) **Social approach.** Representative images indicated the social preference (social and non-social interests) in three chambers of WT and KO mice. Both KO and WT mice spent more time in the mouse-containing chamber (S1) than in an empty cage-containing chamber (E). (B) Social novelty. Representative images indicated the social preference (social novelty) in three chambers of WT and KO mice; the wild-type mice spent more time for a novel mouse-containing chamber (S2) than in a familiar mouse-containing chamber (S1), whereas the KO mice had no preference for either chamber. (C) Reciprocal interaction tests. Social recognition of WT and KO mice was analyzed by measuring the number and time of close interactions. (D) Tube test for social dominance. KO mice experienced significantly more wins than controls. (E) Marble burying test. KO mice buried more marble than controls. (F) Nest building. Representative images indicated the nest quality of WT and KO mice. KO mice had a decreased nest-building index compared with WT mice. (G) Y maze. (a, b) Representative illustrative examples of WT and KO mouse travel pathways on the Y maze. (c) KO mice spent less time on their new arms than wild-type mice. (H) Open field tests. Representative images indicated typical examples of WT and KO mice exploratory behaviors in the 5-min duration open field test (a). KO mice exhibited poor motor ability and spent less time in the center area and increased self-grooming (b). Data information: Mean ± s.e.m. from 10-11 mice per genotype, one-way ANOVA, * P<0.05; ** P<0.01; *** P<0.001).

## Discussion

In the present study, we demonstrated that dcf1 knockout causes delayed neuronal migration, axonal swelling and abnormal boutons. Western blot results showed significant differences in MAP-2 and doublecortin (DCX) expression between KO and Wild-type mice (FIG S1B). MAP-2, a major member of neuronal microtubule-associated proteins (MAPs), has functions in neuronal migration, axonal and dendritic morphogenesis and neurite outgrowth by organizing microtubules (Conde & Caceres, 2009; Teng et al, 2001). Doublecortin (DCX), another MAP enriched at the tip of growing neurites, is thought to play a role in the consolidation phase of axon formation and radial migration (Bielas et al, 2007; Conde & Caceres, 2009). Microtubule is the executor of neuronal migration and morphogenesis, and MAPs regulates assembly, organization and dynamics of microtubule, so the affected MAPs will definitely affect neuronal migration and morphogenesis. DCF1 knockout leads to changes in MAPs expression, which makes it easy to understand why DCF1 deletion affects neuronal migration and axon morphogenesis.

It is traditionally believed that the ERK signal is mainly involved in proliferation, differentiation, learning and memory, development, synaptic plasticity and neural apoptosis (Li et al, 2014; Miningou & Blackwell, 2020). The role of ERK in neuronal migration and autism spectrum disorders (ASD) has long been ignored and has only been reported in recent years (Vithayathil et al, 2018; Xing et al, 2016). In this study, we investigated how the cell behavior of radial migration of cortical neurons is regulated. We found that knockout of dcf1 resulted in delayed neuronal migration and hyperactivated ERK signals. More importantly, we demonstrated that the defects in neuronal migration in dcf1 KO mice can be reversed by inhibiting the ERK signaling pathway. Overexpression of DCF1 led to downregulation of ERK activity, which could not be reversed even by PMA stimulation. PEBP1, a RAF kinase inhibitor that inhibits ERK activation by directly binding RAF1, thereby preventing its phosphorylation (Vandamme et al, 2014; Wu et al, 2014). PEBP1 was involved in the regulation of the ERK signaling pathway in this research. We found biochemically interaction between DCF1 and PEBP1, and the expression level of PEBP1 was affected by DCF1. The co-localization of DCF1, PEBP1 and RAF1 in primary neurons was confirmed. The fact that overexpression of PEBP1 could repair the lesions caused by DCF1 deletion implies that PEBP1 may mediates the regulation of the ERK signaling by DCF1.

The development of neocortex requires precise regulation of neuronal migration, and abnormalities may lead to serious neurological diseases such as ASD(Bocchi et al, 2017; Magnusson et al, 2012; Wegiel et al, 2010), which is characterized by poor social interactions, repetitive behaviors and altered communication(Rapin & Tuchman, 2008). Here we demonstrate that delayed migration is associated with long-term behavioral abnormalities, from ASD-like social deficits to compulsive behavior and cognitive flexibility. Together with previously reported phenotypes of anxiety, learning and memory disorder (Liu et al, 2018), these results indicated that dcf1-/- mice phenotype is all the major symptoms of autism. Although autism is associated with brain development, the mechanism is poorly understood. We hypothesized that DCF1 regulated the ERK signaling pathway through PEBP1, changed the expression of the downstream map, affected the assembly of microtubule, and then affected the migration and morphogenesis of neurons, and changed the neural circuits, leading to autism. These hypotheses demonstrate the critical role of dcf1 in regulating the early development of the cerebral cortex, thereby providing a possible explanation for the genesis of autistic-like behaviors in dcf1-deficient mice.

## Materials and Methods

### Animals

We generated mutant mice that lacked the dcf1 gene via embryonic stem cell-mediated gene targeting as described previously (Liu et al, 2018). The dcf1-/- mice on the C57BL/6J background had a normal appearance compared with the wild-type (WT) littermates. Mice were housed, bred, and treated according to the guidelines approved by the Home Office under the Animal (Scientific Procedures) Act 1986.

### Tissue preparation and histology

All embryos were collected and fixed by immersion in 4% PFA in 0.1M PBS, pH 7.4, for 4h at 4°C. Postnatal mice were perfused with 4% paraformaldehyde in 0.1M PBS. The brain tissues were isolated and further fixed in 4% PFA for 24 h. After fixation, brains were washed with PBS, cryoprotected with 30% sucrose, then frozen in O.C.T. compound (Sakura Finetek) and sliced (20um) with a cryostat (Thermo Fisher scientific). Cryo-sections were air dried at room temperature and stored at −80°C until used.

### Cell culture and transfection

Primary neurons from P0 mouse brains’ cortex were cultured with NEURAL BASAL supplemented with GLUTAMAX and B27, and transfected with Calcium Phosphate. HEK293T were cultured in DMEM supplemented with 10% FBS. Neuro2A (N2A) cells were cultured in MEM supplemented with GLUTAMAX and 10% FBS. All cells maintained in a humidified incubator with 5% CO_2_ at 37 °C. HEK293T and N2A cells were seeded 1 d before transfection at a density that reached 50% confluency on the following day. Plasmids were transfected into cells with Lipofectamine 2000.

### Immunohistochemistry and image analyses

Cells were plated on coverslips pre-coated with poly-L-lysine for 24 hr. Before immunostaining, cells were fixed using 4% paraformaldehyde. Coverslips or cryo-sections were stained with standard protocols by using the following primary antibodies: mouse anti-NEUN (1:500), rabbit anti-MAP-2 (1/1000), goat anti-DCX (1/100), rabbit anti-CUX1 (1/100), rabbit anti-PEBP1 (1:100), mouse anti-raf1 (1/1000), goat anti-GFAP (1:2000), rabbit anti-NEUN (1:500), mouse anti-oligo2 (1/200), mouse anti-NG2 (1:500), goat anti-GFAP (1:2000), rabbit anti-Tuj1 (1:500).Cells or sections were then incubated with appropriate fluorescent secondary antibodies. DNA was stained with 0.5 μg/ml DAPI. Images were acquired on Nikon TI-S inverted microscope, Zeiss LSM 710 confocal microscope.

### Co-Immunoprecipitation and Immunoblotting

For Co-Immunoprecipitation experiments, Cultured cells were lysed in buffer (75 mM HEPES pH 7.5, 150 mM KCl, 1.5 mM MgCl2, 1 mM EGTA, 10% glycerol, and 0.075% NP-40) supplemented with and protease inhibitor cocktail and phosphatase inhibitors. The Co-IP was conducted under the instruction of Dynabeads Protein G (Thermo Fisher scientific, Cat.10003D) protocol. In brief, HEK293T cell was transfected with pcDNA3.1-DCF1-his-myc or pcDNA3.1-PEBP1-1d4. After 44 hours, myc mouse monoclonal antibody, his mouse monoclonal antibody, 1D4 mouse monoclonal antibody and mouse IgG was incubated with Dynabeads Protein G for 3h on a rocking plate at RT separately. 2h later, cell was lysed in RIPA buffer supplemented with protease inhibitor cocktail and phosphatase inhibitors. Then the extracts were incubated with mixture above for 3h on a rocking plate at RT, washed with 50mM Glycine buffer, pH2.8. followed by western blotting with a rabbit polyclonal anti-DCF1 antibody, rabbit polyclonal anti-pebp1 antibody and mouse monoclonal anti-raf1 antibody. For immunoblotting, cultured cells and tissue samples were lysed in RIPA buffer cocktail and boiled in SDS sample buffer. Samples were resolved over 12 or 15% SDS-polyacrylamide gels and transferred onto nitrocellulose membranes. The membrane was probed with primary antibodies. Antibody concentrations were as follows: anti-PEBP1, 1:100; anti-RAF1, 1:1000; anti-GFAP, 1:2000; anti-NEUN, 1:500; anti-MBP, 1:1000; anti-GAPDH, 1:1000; anti-β-actin, 1:1000; anti-JNK, 1:100; anti-p-JNK, 1:100; anti-ERK, 1:100; anti-p-ERK, 1:100; anti-JNK, 1:100. After incubation with secondary antibodies, affinity purified with DyLight 800 or DyLight 680 conjugate, 1:10000. Images were visualized by Odyssey infrared imaging system at 700 nm and 800 nm in 16-bit TIFF format.

### In utero electroporation

Plasmids were transfected by in utero electroporation in accordance with previous methods(Saito, 2006). In brief, multiparous wild type or DCF1 knockout mice at 14.5 d of gestation were anesthetized with 2% sodium pentobarbital (3.3 ml/ kg, intraperitoneally). Uteruses were exposed, and then 8-12 ug of plasmid mixed with Fast Green (2 mg/ml; Sigma) was injected into the lateral ventricle of embryos. Next, electric pulses were generated by an ECM830 (BTX) and applied to the cerebral wall at five repeats of 30 V for 50ms, with an interval of 100ms.

### Slice culture and Time-lapse imaging

Brains transfected with EGFP or DCF1 at E14.5 were dissected from E17.5 mice in ice-cold Hanks’ balanced salt solution (HBSS), embedded in 4% low-melting point agarose in HBSS, and sectioned (coronal) on a vibratome at 100μm. Slices were cultured on Millicell-CM inserts (Millipore) in a 35-mm culture dish containing 1 ml of medium (DMEM plus 10% fetal bovine serum and 2% B27; Gibco) and were incubated in the cell culture incubator for 1 h to allow recovery from injury. The culture dish was then transferred to live cell station on the inversed fluorescent microscope (Zeiss LSM 510). The tissue was illuminated with a 488-nm wavelength light. Images were acquired for four hours.

### Isothermal titration calorimetry

Dcf1 and PEBP1 were cloned into pCold-TF vector and purified through affinity chromatography from expression in E. coliBL21(DE3). Thrombin was used to digest fusion proteins to remove TF. ITC experiments were performed on a MicroCal iTC200 system (GE Healthcare) in 0.01 M PBS pH 8.0 at 25 °C and with constant stirring at 1,000 r.p.m. Prior to experiments, the protein was dialyzed into 0.01 M PBS pH 8.0. The concentration of DCF1 was adjusted to 10μm, and PEBP1 to 100μm. A typical experiment consisted of 20 subsequent injections with a 2μl injection volume into a cell filled with 200μl PEBP1. Each injection was made over a period of 4 s with a 120s interval between subsequent injections. Heat released by each injection was recorded at ‘high’ gain setting, and the background was subtracted. Data analysis was carried out with OriginPro ITC analysis package.

### Behavioral Assays

Three-chamber social preference test. The three-chamber was used to assess social approach and preference for social novelty. The three-chamber apparatus was a Plexiglas box (60 × 45 × 24cm, L × W × H) with two transparent partitions, which respectively constituted the left, center, and right chambers. In the habituation phase of 10min, two empty round wire cages with a diameter of 10 cm) were placed in the left and right chambers respectively, and experimental mouse (3-month-old males) was placed in the central chamber and allowed to freely explore each chamber. Afterwards, an age- and gender-matched unfamiliar stranger mouse (M1) was placed in one of the two wire cages. The other one remained empty (E). The tested mouse was recorded for 10 min. In the last 10-min session, a second stranger mouse (M2) was placed in one wire cage, which previously served as an empty cage. The test mouse was placed in the center, and allowed to freely explore the chamber for 10 min. The movement of the mouse was recorded by a video camera. Video tracking software was used to measure the time spent in each chamber, and within a 5 cm radius proximal to each wire cage. (EthoVision XT 5.0) Reciprocal interaction test. The social interaction in pairs (WT–WT or WT-KO or KO-KO) test was performed in a clear cage (20×45×20cm, L × W × H). Tested mice (3-month-old males) were first habituated for 2 days in 10-min sessions and on day 3 pairs of familiar and unfamiliar mice were placed into a new cage. The social interactions were recorded by video camera for 10 min. The time spent in close contact (indicative of social interaction) and the frequency of close contact were measured manually by a researcher who was blind to the genotype of test mice. Close contact was defined as close huddling, sniffing (nose-to-nose, anogenital sniffing) or all grooming.

Social Dominance Tube Test. The tube test was used to assess social dominance in mice. The tube apparatus (30×3.5 cm, L × D) was made of clear Plexiglas material. It consists of two start areas, a two-section tube and one neutral area between the two-sections. Two age- and gender-matched tested mice (WT-KO) were placed at the opposite ends of the tube and began to explore in a forward direction. At the end of the test, withdraw from the test tube, the score was zero (lost), and the opponent’s score was 1 (win). Each test mouse was challenged three times with a novel strange mouse (a match). Calculate the percentage of each mouse winning in three tests, and then calculate the average winning rate for each group.

#### Marble burying

Marble burying was a repetitive digging behavior and used to assess obsessive-compulsive behavior in mice. Tested mice (3-month-old males) were habituated to clean Plexiglas cages(30×18×12cm) filled with a 4 cm thick layer of clean corncob bedding for 10 min. Afterwards, mice were presented with 15 glass marbles in a 5×3 grid placed on the surface of bedding and allowed 45 min to bury marbles. The number of marbles buried was recorded.

#### Nesting behaviour

Nest building activity was monitored compulsive-like behaviors in both male and female mice. Single-housed mice (3-month-old males) were transferred into a new cage with nest-building material, three 5 × 5 cm square of white compressed cotton pads (Nestlets TM; Ancare, Bellmore, NY) in a random corner. After 24 h, nest building was assessed based on a 5-point scale, 1: Nestlet not noticeably touched; 2: Nestlet partially torn; 3: Nestlet mostly shredded, but no identifiable nest; 4: identifiable, but flat nest; and 5: a near perfect nest.

#### Y-maze

A Y-maze was used for a cognitive flexibility test. A Y-maze was made of gray plexigals, and composed of three equal-sized transparent arms (30 × 9× 16cm, L × W × H) with removable gates. The three identical arms were randomly designated: start arm, in which the mouse started to explore (always open), novel arm, which was blocked during the 1st trial, but open during the 2nd trial, and other arm (always open). During the first 10-min trial, the tested mouse was only allowed to explore two arms (stat arm and other arm) by closing the gate of the novel arm. For the second trail, during which all three arms were accessible, mouse was placed back in the same stating arm, with free movement for 5 min. The movement of the mouse was recorded by video camera. Time spent in each arm was further analyzed by video tracking software (EthoVision XT 5.0, Noldus Technology).

#### Open field test

The open field test is a measure of assessing mice tendency to explore novel objects and environment. Tested mice (3-month-old males) were placed near the wall-side of the novel large open arena (70cm × 70cm) and allowed to explore the new environment for a total of 10 min. Mouse movements were recorded. The time spent and frequency in the center (35 × 35 cm) and total distance was analyzed. (EthoVision XT 5.0) Time of self-grooming behavior was analyzed.

### Imaging and Statistics

Images were processed and analyzed using image pro plus 6.0 software. Statistical analysis and graph creation were performed by Excel and SPSS software. Results were presented as means ± SEM. Student’s t-test and two-way ANOVA was used for comparisons between dcf1−/− mice and their wild-type controls.

## Acknowledgements

This work was funded by the National Science Foundation of China (81271253, and 81471162), the Science and Technology Commission of Shanghai (14JC1402400).,.

## Author contributions

R.L.F. and T.Q.W. conceived of the project and designed the experiments and analyses. R.L.F. carried out the experiments and the analyses. Y.L.C. and J.J.G. executed in utero electroporation. Y.Y.S did the ITC experiment, G.H.L. performed the Co-Immunoprecipitation experiments. Q.L. and X.C.J. constructed the plasmid pCAGGS-DCF1-EGFP and N2-Dcf1-EGFP. J.W. generated the knockout mice Dcf1-/-. R.L.F. and T.Q.W. wrote the manuscript.

## Expanded View Figure legends

**Fig S1.**
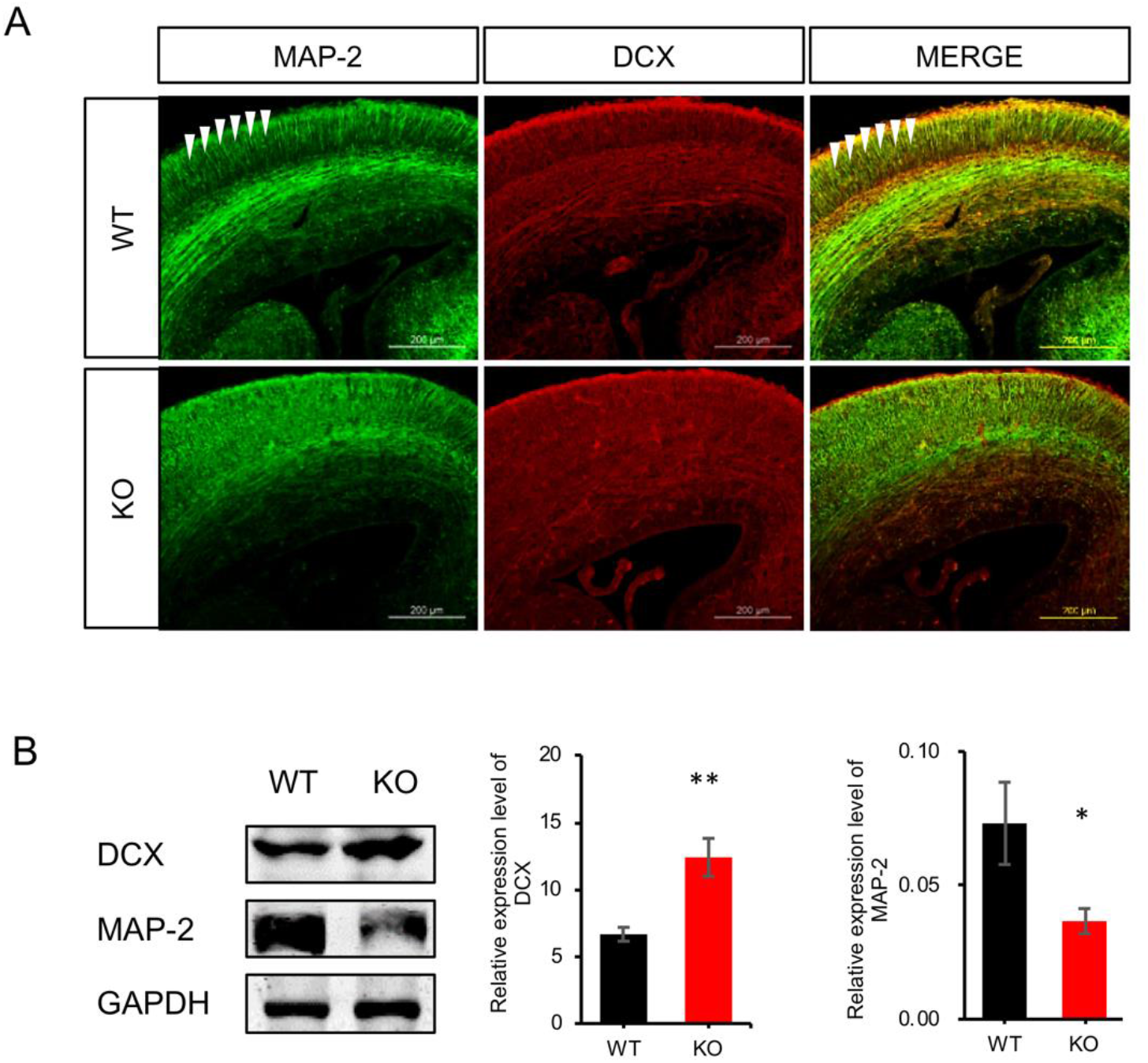
Dcf1 is required for a cortical radial pattern. (A) Anti-MAP-2 and anti-DCX antibodies were used to visualize neurons in coronal sections at E17.5. In the WT mice, cortical neurons were tightly clustered in a radial pattern as indicated by arrowhead, whereas in the *dcf1*-/- mice, the cortical neurons were disorganized with no clear radial pattern. (B) Expression levels of MAP-2 and DCX in the cortex were analyzed by western blot at E17.5, and the differences between KO and WT mice were compared by statistical analysis.

**Fig S2.**
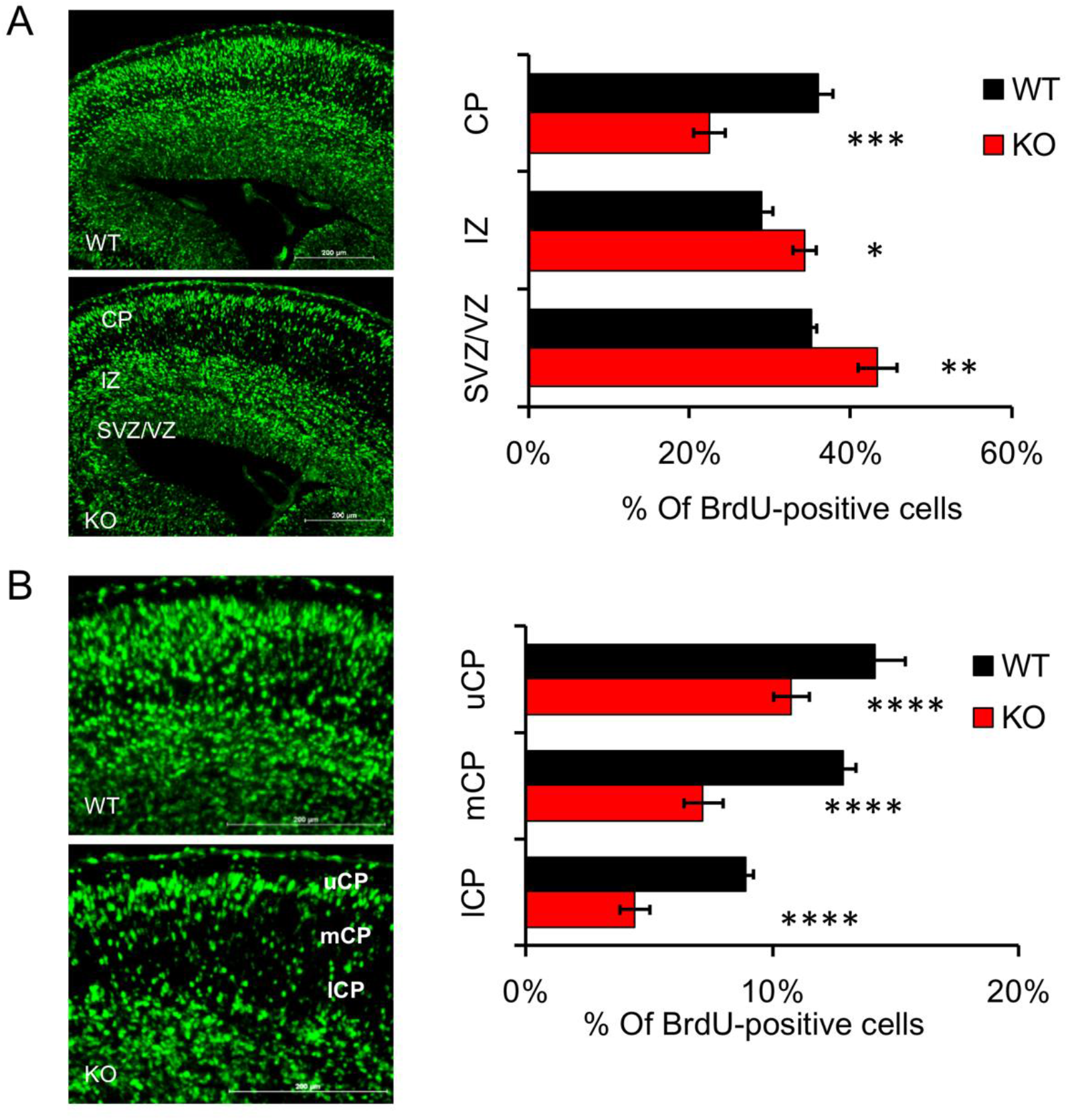
BrdU-labeling pattern in the brains of *dcf1-/-* mice and their WT littermates. Dcf1 heterozygous mice were mated with heterozygous mice, pregnant mice were injected with BrdU at E14.5. Embryos were examined at E17.5. (A) BrdU-labeled cells of dcf1-null mice were significantly decreased in CP and increased in IZ and VZ/SVZ compared with their littermate controls. (B) Quantitative analysis distribution of BrdU^+^ cells in CP. Compared with littermates, the number of BrdU-positive cells in each layer of the CP was dramatically reduced in dcf1-/- mice. (CP, cortical plate; uCP, upper CP; mCP, median CP; lCP, lCP; IZ, intermediate zone; SVZ/VZ, subventricular zone/ventricular zone). Data was shown as mean ± sem, Student’s t test; *p < 0.05, **p<0.01, ***p<0.001, ****p<0.0001.

**Fig S3.**
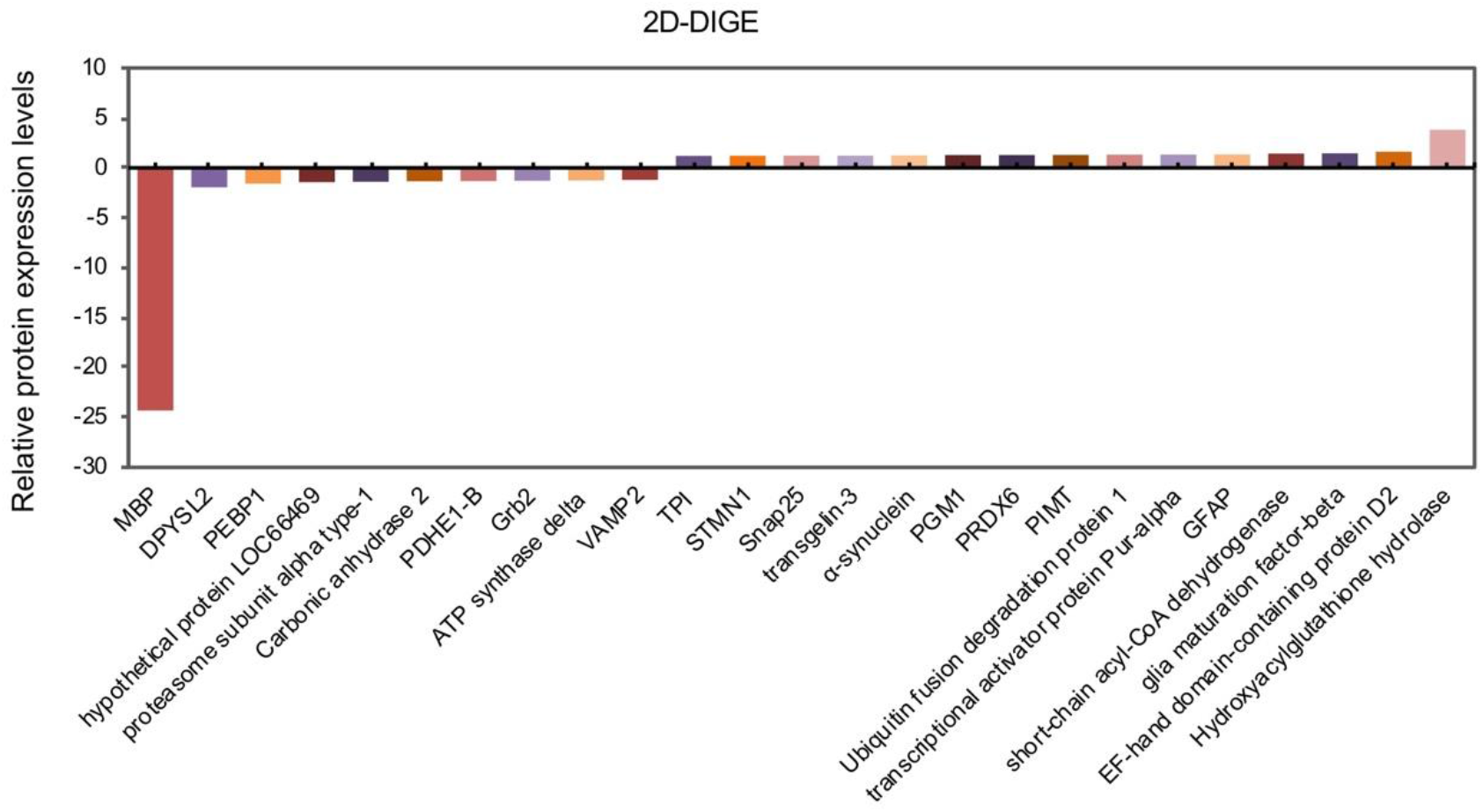
Identification of Differentially Expressed Proteins using 2D-DIGE and Mass Spectrometry. Using 2D-DIGE and mass spectrometry, 25 differentially expressed proteins (fold change>1.5, P<0.05) were identified in the adult cortex of dcf1-/- and WT mice.

**Supplementary Movie:** Real-time imaging of cultured slices was obtained from sections of E17.5 brains electroporated at E14.5. Movies indicate the migration of neurons in the slices during a 4-h incubation on a Live Cell Station, **(Movie s1)** Wild type mouse electroporated with pCAGGS-EGFP; **(Movie s2)** *Dcf1*-/- brains electroporated with pCAGGS-EGFP; **(Movie s3)** *Dcf1*-/- brains rescued using pCAGGS-DCF1-EGFP.

